# Subcellular proteomics of *Paramecium tetraurelia* reveals mosaic localization of glycolysis and gluconeogenesis

**DOI:** 10.1101/2025.04.24.650466

**Authors:** Dagmar Jirsová, Timothy J. Licknack, Yu-Ping Poh, Yanping Qiu, Natalie Quan, Tsui-Fen Chou, Tim Karr, Michael Lynch, Jeremy G. Wideman

**Author notes:** These authors contributed equally and their surnames are in alphabetical order (Dagmar Jirsová, Timothy J. Licknack, Yu-Ping Poh).

## Abstract

Ciliates are unicellular heterotrophic eukaryotes, most of which consume other microbes as prey. They exhibit nuclear dimorphism which requires reconstruction of a transcriptionally active macronucleus from the germline micronucleus after sexual recombination. This complex genomic structure has prevented the development of highly tractable genetic models leaving much of ciliate cell biology unexplored. To complicate matters further, some ciliates tend to accumulate many gene duplicates either singly or via whole genome duplications. Thus, extensive insight into the cell biology of ciliates requires the use of high-throughput tools like subcellular proteomics.

Here, we use a subcellular proteomics workflow to classify over 9,000 proteins to 16 subcellular compartments in *Paramecium tetraurelia*. From these data, we identify a small but robust subcellular cluster containing canonical mitochondrial outer membrane proteins as well as some ER proteins, putatively at membrane contact sites. Within this cluster, we identified the important glycolytic enzyme phosphofructokinase, which contained a transmembrane domain. Further investigation revealed that several latter-acting glycolytic enzymes were localized to the mitochondrial cluster. The location of phosphoenol pyruvate carboxykinase and pyruvate carboxylase in the mitochondria but pyruvate kinase in the cytosol suggests that ciliates prefer gluconeogenesis over glycolysis. The localization of these enzymes was confirmed in a preliminary subcellular proteome of *Tetrahymena thermophila*. In sum, our findings suggest that mitochondrial localization of glycolytic/gluconeogenic enzymes is widespread across ciliates and that several may preferentially undergo gluconeogenesis over glycolysis using amino acids as a primary carbon source in both catabolic and anabolic metabolism.

**Highlights:**

Subcellular proteomics of *Paramecium tetraurelia* revealed that glycolytic and gluconeogenic enzymes are mosaically distributed between the cytosol, mitochondrial matrix, and mitochondrial outer membrane.
A distinct mitochondrial outer membrane compartment was identified with 105 classified proteins, including core mitochondrial biogenesis proteins and a putative Tom70-like protein.
Phosphofructokinase, a key glycolytic enzyme, was found embedded in the mitochondrial outer membrane.
Localization of biochemical pathways suggest ciliates favor gluconeogenesis over glycolysis.
In total, over 9000 *Paramecium* proteins were identified using subcellular proteomics and classified into 16 different cellular compartments.

## Introduction

Ciliates are among the most complex and diverse cells of any lineage of unicellular eukaryotes ^1^. Some species are as large as some animals, and many have extremely complex cells. Their name comes from the arrays of coordinated cilia, which, in many species, cover almost the entire cell body. They exhibit a characteristic form of nuclear dimorphism, with somatic macronuclei and germline micronuclei. Though they are probably the most thoroughly studied group of (mostly) free-living heterotrophic protists, relatively little is known about the protein contents of their subcellular compartments (e.g., ^2^). Previous work has suggested that some ciliates can have a rather peculiar localization of biochemical pathways, such as localization of some glycolytic enzymes to the mitochondria. For instance, phosphofructokinase activity was seen on isolated *Tetrahymena pyriformis* mitochondria ^3^, which was later confirmed and expanded to five other glycolytic enzymes via mass spectrometry of isolated mitochondria in *Tetrahymena thermophila* (*Tetrahymena* henceforth) ^4^. Whether this is a specific feature of *Tetrahymena* or if it can be generalized to all ciliates remains unclear.

Non-cytosolic localization of glycolytic enzymes is not unique to ciliates and has been observed in a couple of other eukaryotic lineages ^5^. For example, in trypanosomes and other kinetoplastids, the first seven steps of glycolysis are localized to modified peroxisomes called glycosomes ^6,7^. Glycosomal localization of glycolysis has also been inferred in the lineage sister to kinetoplastids, the diplonemids ^8^, though data are limited ^9^. In stramenopiles like diatoms and oomycetes, ATP-yielding steps of glycolysis are localized to both mitochondria and the cytosol. However, in another stramenopile, the human intestinal commensal *Blastocystis,* these enzymatic reactions localize exlusively to mitochondria ^10^. Similarly, the parasitic rhizarian *Paramikrocytos canceri* localizes a similar set of enzymes to its extremely reduced mitochondria-related organelle ^11^. Though it is unclear why different lineages localize glycolysis to different cellular compartments, it is possible that compartmentalization of glycolytic enzymes might offer some selective advantages. For example, a selective advantage could be gained by separating reactions that require different conditions (e.g., pH, or specific reactant/product concentrations) thereby improving the overall efficiency of the cell. However, the case in ciliates remains unclear, and further investigation into diverse ciliates is necessary to answer this question.

One of the most famous and best studied ciliates is *Paramecium tetraurelia* ^12,13^ (*Paramecium* henceforth). *Paramecium* contains about 44,000 protein coding genes which resulted from three ancient, whole-genome duplications ^14,15^. Despite being a ciliate “flagship” our knowledge about its cell biology is limited to only a few hundred proteins that have been localized within the cell or functionally investigated ^2^. Genetic manipulation of *Paramecium* is difficult and time-consuming ^16^. Therefore, high-throughput proteomics methods are needed to bridge the gap and provide extensive information about protein localization and function this organism.

Subcellular proteomics methods ^17^ enable the localization of thousands of proteins to cellular compartments in a single experiment and the generation of subcellular protein atlases ^18–24^. These various techniques stand out as they can even be used to investigate non-model systems or organisms that are difficult to genetically manipulate (e.g., ^25^). Thus, these methods provide opportunities for high-throughput identification of organellar components across diverse and complex cells, including ciliates like *Paramecium*.

Here, we apply a subcellular proteomics workflow to *Paramecium* and classify over 9000 proteins to 16 cellular compartments to produce the first ciliate subcellular atlas. We confidently predict 1,089 proteins to the mitochondria and 353 to the cytoplasm. We also identified a small but robust cluster of 42 mitochondrial outer membrane proteins (MOM). Our analyses demonstrate that glycolytic enzymes are mosaically distributed between the cytosol and mitochondria, with phosphofructokinase specifically embedded in the MOM. These *Paramecium* data are confirmed in a preliminary subcellular atlas of *Tetrahymena*. Because the last step of glycolysis, which produces pyruvate is located outside the mitochondria in the cytosol, we explored the possibility that ciliates get most of their energy from digesting amino acids and preferentially undergo gluconeogenesis rather than glycolysis. Comparative analyses suggest that these findings are generalizable to diverse ciliates.

## Results and Discussion

### Subcellular proteomics leads to confident prediction of over 4500 proteins to subcellular compartments in *Paramecium tetraurelia*

To determine the subcellular location of thousands of *Paramecium* proteins, we used a modified subcellular proteomics workflow (Figure 1A) adapted from previous studies ^21^. For each experiment, between two and ten million cells were harvested and lysed using a nitrogen cavitation chamber (see methods). Cell lysates were then subjected to differential centrifugation resulting in 11 distinct fractions (Figure 1A), plus a macronucleus-enriched fraction ^26^. Immunoblotting was used to confirm distinctness of fractions (Figure S1). All fractions were then subjected to label-free quantification (LFQ) mass spectrometry using an Orbitrap Fusion Lumos Tribrid Mass Spectrometer (see methods). We detected a total of 9,026 unique proteins in our dataset (Table S1). After normalization and imputation, relative protein abundance profiles were plotted for each protein (e.g., Figure 1B). The final, normalized, and imputed data were analyzed and visualized as a t-distributed stochastic neighbor embedding plot (t-sne) using the pRoloc R package ^27^ (Figure 1C).

**Figure 1.**
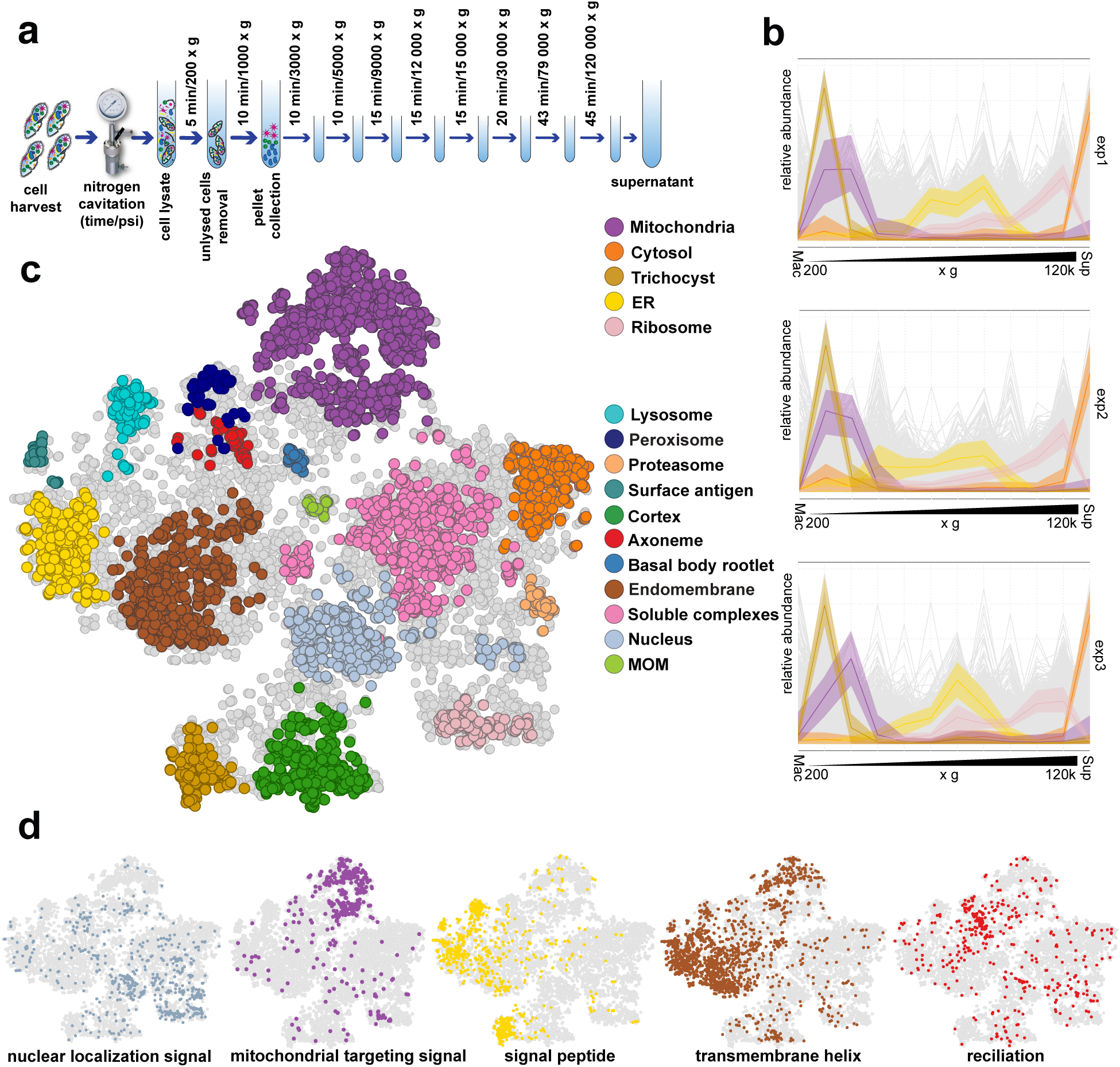
Subcellular proteomics of *Paramecium tetraurelia*. (a) Subcellular proteomics workflow. We used nitrogen cavitation to lyse harvested cells and followed the differential centrifugation protocol published by Geladaki, et al. (2019). (b) Protein abundance profiles of mitochondrial, cytosol, trichocyst, ER, and ribosomal clusters. After label-free quantification mass spectrometry, data processing, and mapping of marker proteins, we created abundance-distribution profiles of selected subcellular marker proteins measured in each experiment. Individual fractions contained different enrichments of organellar proteins. (c) Subcellular protein atlas of *Paramecium tetraurelia*. We used t-SNE projection to visualize the distribution of organellar proteins after SVM classification. (d) Protein targeting signals align overlap with their predicted organelles in (c). Using available prediction programs, we predicted nuclear localization signals, mitochondrial targeting signals, signal peptides, and transmembrane helixes onto our subcellular atlas. We also mapped a publicly available transcriptome of reciliation onto the subcellular atlas.

To determine the relative completeness of our detected proteome, we computed Benchmarking Universal Single-Copy Orthologs (BUSCO) completeness scores ^28,29^. Our proteomic dataset had a relatively high alveolate BUSCO score (93.6%), which is comparable to the score of the complete *Paramecium* genome (98.8%), though we only detected ∼ ¼ of the predicted *Paramecium* proteome (e.g., 9,026 of > 44,000). The relatively high BUSCO is likely explained by the fact that many BUSCO proteins are highly expressed and therefore highly abundant and more readily detected by mass spectrometry. The similarity between detected and total proteome BUSCO scores is also seen in the organellar proteome of *T. gondii* ^22^ in which 52% and 61% of core eukaryotic proteins were identified in their organellar and predicted proteomes, respectively. The 5-9% of missing BUSCOs could represent genes that are conserved but have low expression or are highly hydrophobic—both qualities that make proteins difficult to detect by mass spectrometry ^30^.

To predict the cellular location of thousands of proteins, we used a 291-protein marker set with known localizations (see Figure S2A and Table S1). We used these marker proteins to perform supervised classification on all 9,026 proteins using a support-vector machine (SVM) classifier ^31^ (Figure S2B) and chose a median cutoff resulting 4,513 predicted proteins across 16 cellular compartments (Figure 1C). The median cut-off did not bisect each compartment evenly, as some had much higher median SVM scores (e.g., ribosomes and mitochondria) than others (e.g., axonemes), likely the result of some compartments forming tighter clusters than others. When we refer to *predicted* proteins, we are referring to proteins that fall within the median cutoff (i.e., colored proteins in Figure 1C). When we refer to proteins as *classified* to particular compartments, we refer to all proteins classified to a single compartment (Figure S2B). To determine how well our biologically informed SVM clusters align with clusters generated *de novo*, we generated 16 clusters using the k-Means (KM) algorithm ^32^ (Figure S2C). Qualitatively, many KM clusters overlapped well with organellar predictions (Figure S2D).

To further corroborate the validity of our data, we predicted the presence of targeting signals and transmembrane domains using SignalP ^33^, NLStradamus ^34^, MitoFates 1.2 ^35^, and DeepTMHmm 1.0.24 ^36^ (Figure 1D). Nuclear localization signals were present mainly in the nuclear compartment and the ribosome. Predicted mitochondrial targeting signals were enriched in the mitochondrial compartment. As expected, signal peptides were enriched in ER-derived membrane trafficking compartments including the ER, lysosome, endomembrane, surface antigen, and trichocyst compartments. Transmembrane domains were enriched in membrane bound compartments like the mitochondrial outer membrane, ER, and endomembrane compartments but largely absent from other compartments. We used previous data ^37^ to map proteins expressed during reciliation showing that axonemal proteins are enriched in this dataset. These data demonstrate that predicted compartments are enriched for proteins that are predicted to function or reside in these compartments strongly suggesting that our data can be used for functional inference and as a hypothesis-generating tool for downstream investigations.

### Subcellular proteomics resolves many cellular compartments in *Paramecium*

Subcellular proteomics of *Paramecium* resolved several large protein complexes and endomembrane compartments. Large cytosolic complexes like the ribosome (167 total proteins) and the proteasome (65 total proteins) resolved well in our data (Figure 1C). We defined several other clusters in our dataset. These included the nucleus (413 proteins), ER (325 proteins), endomembranes (569 proteins), soluble protein complexes (512 proteins), cortex (393 proteins), axoneme (49 proteins), basal body rootlet (28 proteins), trichocyst (206 proteins), and surface antigen (57 proteins) compartments. The axonemal compartment overlapped with a cluster of peroxisomal proteins (68 proteins) which contains many core peroxisomal enzymes as well as membrane-bound PEX proteins. Finally, a tight cluster of lysosomal proteins (142 proteins), many of which contain signal peptides (Figure 1D) was also identified. Follow-up studies will be focused on disentangling the complexities of these various cortical and endomembrane compartments.

Two well-resolved compartments in this study were the mitochondria and cytoplasm (Figure 1C purple and orange, respectively). The cytoplasm contained 343 predicted proteins that largely lack predicted TMDs and contained many canonically cytoplasmic enzymes like aldolase, alcohol dehydrogenase, and various tRNA ligases and synthetases (Table S1). The mitochondrial cluster was the largest of our compartments, containing 1,138 predicted proteins. Our mitochondrial compartment includes all enzymes of the TCA cycle and all electron transport chain complexes except Complex III (likely due to extreme hydrophobicity). In our data, mitochondrial outer membrane (MOM) proteins (e.g., Tom40, Tom22) composed an independent MOM compartment (Figure 2A, light green) with 37 predicted (104 classified) proteins, which exhibited different abundance profiles than all other mitochondrial markers (Figure 2B).

**Figure 2.**
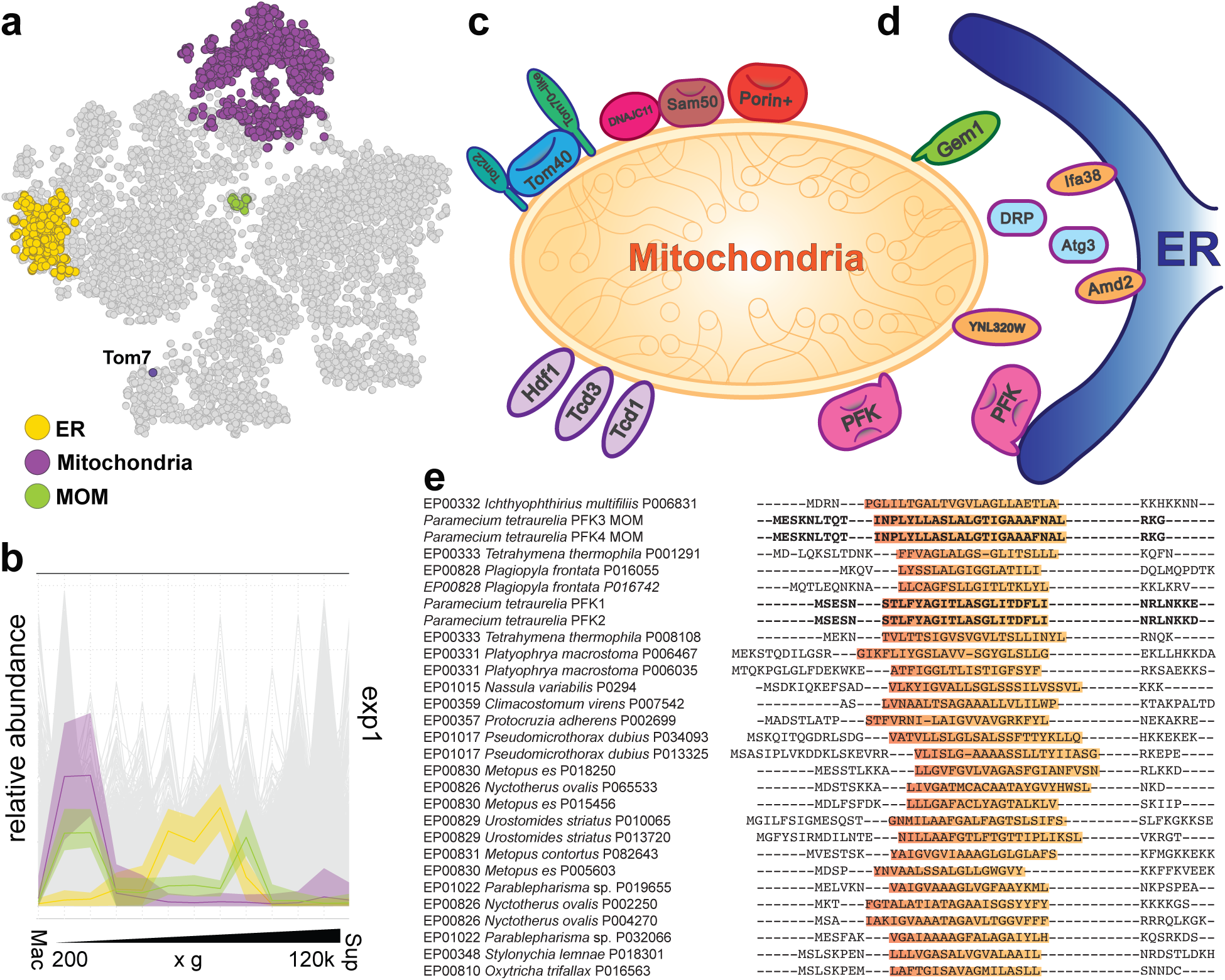
The MOM cluster contained two phosphofructokinase homologues in addition to classic MOM proteins. (a) The mitochondrial outer membrane formed a cluster separate from both the ER and mitochondrial clusters. Tom7 was likely erroneously classified to the trichocyst. (b) The mitochondrial, ER, and MOM clusters have distinct protein abundance profiles. (c) The *Paramecium* MOM contains many classical proteins as well as several putative membrane contact site proteins (d). We identified commonly described MOM proteins including Sam50, DNAJC11, Tom40, Tom22, porin, Gem1, Hfd1, Tcd1, Tcd3, and Amd2. We identified several proteins that act at ER-mitochondria contact sites including dynamin, Atg3, Ifa38, and an orthologue of YNL320W. (d) Ciliate phosphofructokinases (PFKs) have N-terminal transmembrane domains. The *Paramecium* MOM cluster also contains two copies of phosphofructokinase, one of the key regulatory proteins in glycolysis. All PFK proteins obtained from ciliates have a transmembrane domain, suggesting that PFK might be a regular MOM protein in all ciliates.

### The *Paramecium tetraurelia* mitochondrial outer membrane contains much of the expected conserved repertoire

Mitochondrial outer membranes (MOMs) facilitate necessary functions like mitochondrial biogenesis, mitochondrial fusion and fission, organelle communication via contact sites (e.g., with the endoplasmic reticulum), and mitochondrial motility via connections with the cytoskeleton ^38^. Apart from these inevitable MOM functions, a number of enzymes from various pathways are housed at the MOM (e.g., ^39–41^), though the extent to which these auxiliary enzymes have conserved MOM localization across diverse eukaryotes is unclear. In the *Paramecium* MOM cluster, we identified many homologues of the mitochondrial protein import machinery, including the receptor Tom22, and the β-barrels Tom40 and Sam50 (Figure 2C), though Tom7 was absent and instead placed in the trichocyst cluster, likely a rare artifact of imputation. We identified a clear orthologue of the β-barrel Porin as well as a ciliate-specific β-barrel protein similar to Tom40/Porin (Table S1). In addition to these proteins conserved in yeast, we identified an Hsp40-like protein DNAJC11 that is known to interact with the SAM and MICOS complexes in humans ^42,43^. Although, DNAJC11 has been shown to be present across almost all eukaryotes and is likely involved in mitochondrial biogenesis ^44^, its mechanism of function is unclear. We also detected a tail-anchored protein with a TPR-repeat that, when used as a BLAST query into the yeast proteome, retrieves Tom70 as its best hit. Though it is unlikely that this Tom70-like protein is a true orthologue of yeast Tom70, there is precedence of receptor convergence in several other lineages of eukaryotes ^45–48^. Besides this assortment of biogenesis proteins, we detected a few other conserved MOM proteins.

Included in the MOM fraction were several other known proteins involved in mitochondrial division, mitophagy, or cell communication. First, is the enigmatic mitochondrial Rho GTPase (called Gem1 in yeast ^49^ or MIRO in animals ^50,51^). Though nearly universally conserved in the MOM across eukaryotes ^52^, the function of Gem1 remains unclear outside animals, where it is largely known to function in mitochondrial motility in neurons ^51^. Also in the MOM cluster, we identified a ciliate-specific dynamin-related protein (Figure 2D and S3) previously shown to be involved in mitochondrial division ^53^. Next, we identified a homologue of Atg3, an E2 ubiquitin ligase important for the regulation of mitophagy in animals ^54,55^ and whose homologue is involved in mitochondrial homeostasis in the apicomplexan *Toxoplasma gondii* ^56^. In addition to the above-described conserved MOM components, we identified a number of possibly conserved MOM proteins: e.g., orthologues of Tcd1/2, Tcd3, and Hfd1 in yeast ^41^, an orthologue of Amd2 in *Neurospora crassa* ^40^, and an orthologue of the yeast protein YNL320W whose orthologue is present at the MOM in animals ^57^ (see Figure 2D and Table S1). We also observed a number of alveolate, ciliate, or *Paramecium*-specific proteins or paralogues with no obvious homologues in other lineages. Finally, 14 proteins in this cluster contained predicted signal peptides (Table S1), indicating that in addition to MOM proteins, this cluster may also contain ER proteins present at ER-mitochondria contact sites (see ^58^ for a general review on membrane contact sites (MCSs)). In sum, we conclude that the MOM cluster is a robust set of classic mitochondrial outer membrane proteins (e.g., TOM and SAM), their interactors (e.g., Drps and Atg3), a subset of ER proteins likely found at MCSs, and lineage-specific MOM or MCS components. In addition to the various MOM proteins already mentioned, we identified two homologues of phosphofructokinase (PFK), a highly regulated glycolytic enzyme (e.g., ^59^). Both enzymes contain conserved predicted N-terminal transmembrane domains (Figure 2E), both of which have weakly predicted signal peptides indicating possible ER localization at the MCS.

### Glycolysis/gluconeogenesis is localized partly in the mitochondria and partly in the cytoplasm in ciliates

Because PFK and other glycolytic enzymes had been previously detected by biochemical analysis and mass spectrometry in the mitochondria of *Tetrahymena* ^3,4^, we sought to determine whether these glycolytic enzymes could be identified in the *Paramecium* mitochondrial cluster of our dataset. Indeed, our *Paramecium* data shows that glycolysis is mosaically localized to the mitochondrial matrix, the MOM, and the cytosol (Figure 3A). We further confirmed previous data from *Tetrahymena*, by performing a preliminary subcellular proteomics analysis (Figure S4 and S5). The mosaic localization of glycolytic enzymes to mitochondria and cytosol are largely consistent between these two ciliate datasets.

**Figure 3.**
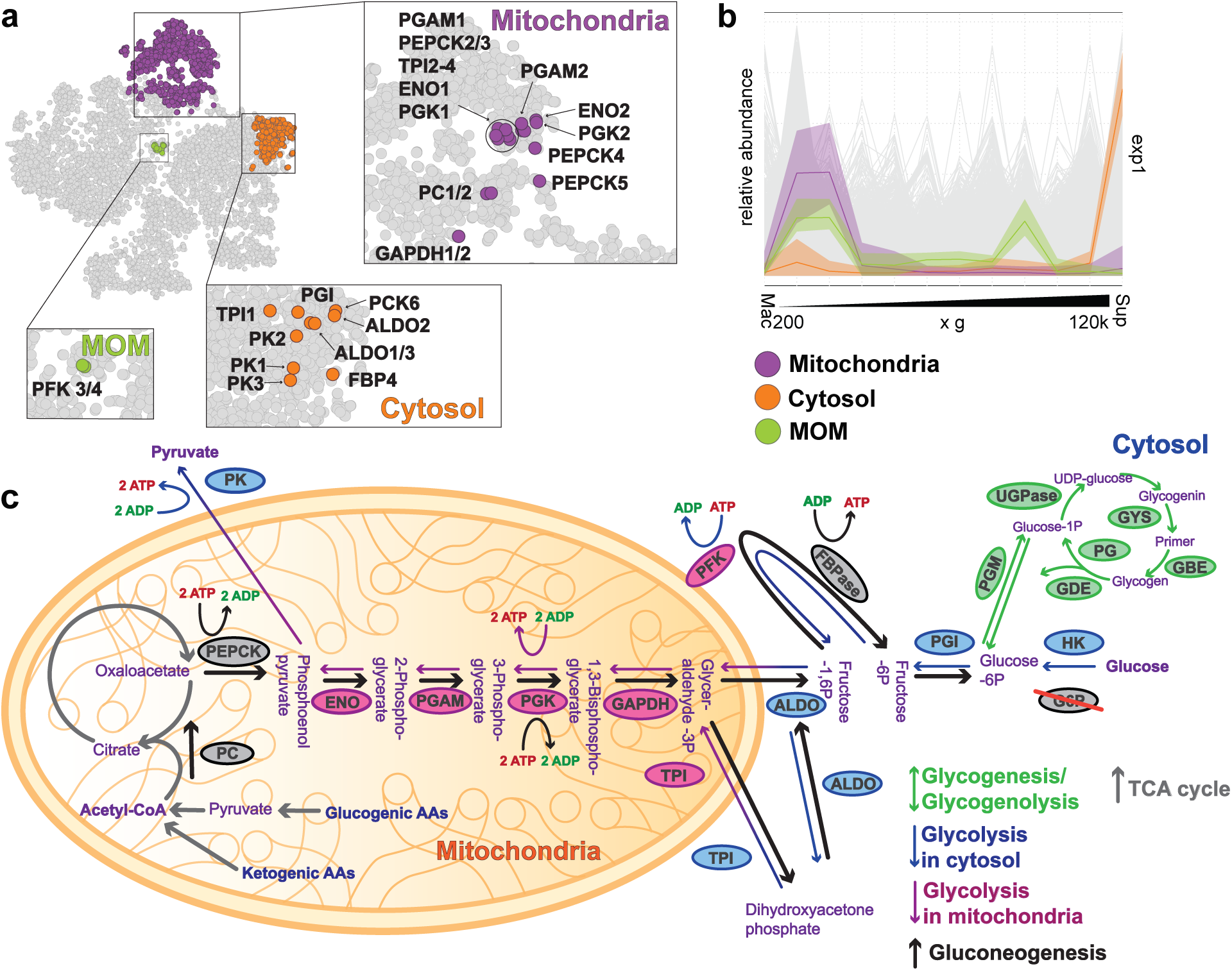
Glycolytic and gluconeogenic proteins exhibit a mosaic localization. (a) Glycolytic and gluconeogenic proteins are distributed between the MOM, cytosol, and mitochondrial fractions. Individual proteins are highlighted in the insets. (b) Mitochondrial, cytosol, and MOM fractions have distinct protein abundance profiles. (c) The first six or seven enzymes that catalyze initial steps of gluconeogenesis are in the mitochondria. The enzymes catalyzing the final steps of gluconeogenesis were found in the cytoplasm. Glycolytic enzymes followed the same pattern (but in reverse), except pyruvate kinase (PK), which was found in the cytoplasm. With this discordance, we asserted that Paramecium prefers gluconeogenesis to glycolysis, storing glucose in glycogen granules.

As an overview, the first steps in glycolysis are uncontroversially located in the cytosol in our datasets (Figure 3B, blue ovals to right), whereas most of the 3-carbon reactions occur in the mitochondrial matrix (Figure 3B, pink ovals), except the final reaction catalyzed by pyruvate kinase (PK), which occurs back in the cytosol. These findings match the data originally produced in *Tetrahymena* ^3,4^, as well as our preliminary *Tetrahymena* subcellular proteomics data (Figure S4). In *Paramecium*, glycolysis begins in the cytosol, where a low-affinity glucokinase (hexokinase IV) adds a phosphate group to glucose, forming glucose-6-phosphate (G6P) (Figure 3C). The next step, catalyzed by phosphoglucose isomerase (PGI), converts G6P to fructose-6-phosphate (F6P), which also occurs in the cytosol. F6P is then phosphorylated by PFKs embedded in the mitochondrial outer membrane (MOM), producing fructose-1,6-bisphosphate (F16P). Aldolase then cleaves F16P into dihydroxyacetone phosphate (DHP) and glyceraldehyde-3-phosphate (G3P). The following step involves triose phosphate isomerase (TPI), where paralogues are found in both the cytosol and mitochondria of *Paramecium*. In contrast, only a mitochondrial form of TPI is present in *Tetrahymena*, suggesting that transporters for G3P and DHP must exist in the mitochondrial membranes of both ciliates. However, transporters specific to these 3C-metabolites have only been experimentally validated in stramenopiles ^60^.

For the subsequent glycolytic steps, including those catalyzed by glyceraldehyde-3-phosphate dehydrogenase (GAPDH), phosphoglycerate kinase (PGK), phosphoglycerate mutase (PGAM), and enolase (ENO), the enzymes are exclusively localized in the mitochondria of *Paramecium* (Figure 3B). In *Tetrahymena*, paralogues of PGAM and ENO are also present in the cytosol (Figure S4). Finally, in both ciliates, the last step of glycolysis, catalyzed by pyruvate kinase (PK), is almost exclusively outside the mitochondria, with one copy potentially dual-localized in *Paramecium* (see below). A few additional paralogues of glycolytic enzymes may be localized to other compartments, though the function of these is unclear (see below). Because compartmentalization is likely adaptive, if glycolysis were running predominantly in the forward direction, one would expect PK, which produces ATP to be localized to mitochondria. We therefore hypothesized that ciliates preferentially use glycolytic enzymes in reverse, for the synthesis of glucose via gluconeogenesis.

### Ciliates prefer gluconeogenesis over glycolysis

Gluconeogenesis can be thought of simply as glycolysis in reverse (Figure 3B). It is the process by which pyruvate is converted into glucose via an 11-step process and functions as a key interface between amino acid, lipid, and sugar metabolism. Only gluconeogenic precursors, that is, certain amino acids and TCA cycle intermediates, can be converted to pyruvate in such a way that glucose can be synthesized without a net loss of carbon. Though pyruvate is not a freely available resource in a ciliate diet, many amino acids (e.g., serine, glycine, alanine, and cysteine) released from the breakdown of proteins from ingested bacteria can be easily converted into pyruvate. Most amino acid metabolic pathways are found in the mitochondria and feed into the tricarboxylic acid cycle (TCA) of most organisms ^61^, which is true also in ciliates (Figure S6). Thus, one starting path for gluconeogenesis begins in the mitochondria with amino acids being converted into pyruvate in the mitochondria. From here, two enzymes, pyruvate carboxylase (PC) and phosphoenol pyruvate carboxykinase (PEPCK) are required to convert pyruvate to phosphoenol pyruvate via oxaloacetate. PC is generally present in the mitochondria of animals ^62^, but is so far exclusively in the cytoplasm of fungi ^63^. PEPCK is found in both the mitochondria and cytoplasm of vertebrates ^64^ but is putatively found only in the cytosol in fungi like yeast. In ciliates, both PC and PEPCK appear to be primarily localized to the mitochondria (Figure 3AB), similar to the location in another alveolate, *Toxoplasma* ^65^. After conversion to PEP, the next five steps catalyzed by ENO, PGAM, PGK, GAPDH, and TPI could all occur in the mitochondria. Catalysis of G3P and DHP to form F16P by aldolase occurs in the cytosol. From here, F16P can be converted to F6P via the gluconeogenesis-specific enzyme fructose 1,6-bisphosphatase (FBP). PGI can then convert F6P to G6P.

Here, the gluconeogenic pathway ends in both ciliates. The final step to form glucose catalyzed by glucose-6-phospate phosphatase is not present in either *Paramecium* or *Tetrahymena*. Instead, glycogenesis occurs via conversion of G6P to glucose-1-phosphate for eventual storage as glycogen ^66,67^. The initial steps of glycogen synthesis are localized to the cytosol in both ciliates, with latter steps either not detected or in a possible endomembrane fraction (Figure S6CD). To determine whether ciliates can grow in media lacking sugars, we compared the growth of *Tetrahymena* in modified Neff media (control), glucose-free modified Neff media, high protein SPP media, and in media with low protein and high glucose concentration (see methods section). *Tetrahymena* grew at approximately the same rate in the presence or absence of sugar (Figure S7). These data suggest that *Paramecium*, *Tetrahymena*, and potentially other heterotrophic ciliates, produce glucose from non-carbohydrate sources, perhaps favoring the anabolic pathway of gluconeogenesis.

### Dual localization of enzymes involved in glycolysis and/or gluconeogenesis in ciliates

Some reactions of the *Paramecium* and *Tetrahymena* glycolytic and gluconeogenic pathways were localized to multiple locations in the cell. In a couple of cases, the multiple localizations were due to paralogues resulting from gene duplications being localized to different compartments. For example, TPI has paralogues clearly localized to either the mitochondria or cytosol in *Paramecium* (Figure 3A); whereas in *Tetrahymena*, PGAM and ENO have paralogues that are differentially localized (Figure S4). Interestingly, a few paralogues of *Paramecium* proteins (PEPCK1, HK1, and PK5) were not clearly localized to any compartment in our analysis (Figure 4A) and instead occupied a space in the t-sne plot between the cytosol and mitochondria clusters (Figure 4A, inset). Though spatial organization on t-sne plots cannot be inferred to have any biological meaning, it can provide hints. On closer inspection, all three proteins had abundance profiles that could indicate that these are dual-localized proteins (Figure 4B). Our data prompted us to take a closer look at the N-termini of these proteins for possible indications of cryptic mitochondrial targeting signals.

**Figure 4.**
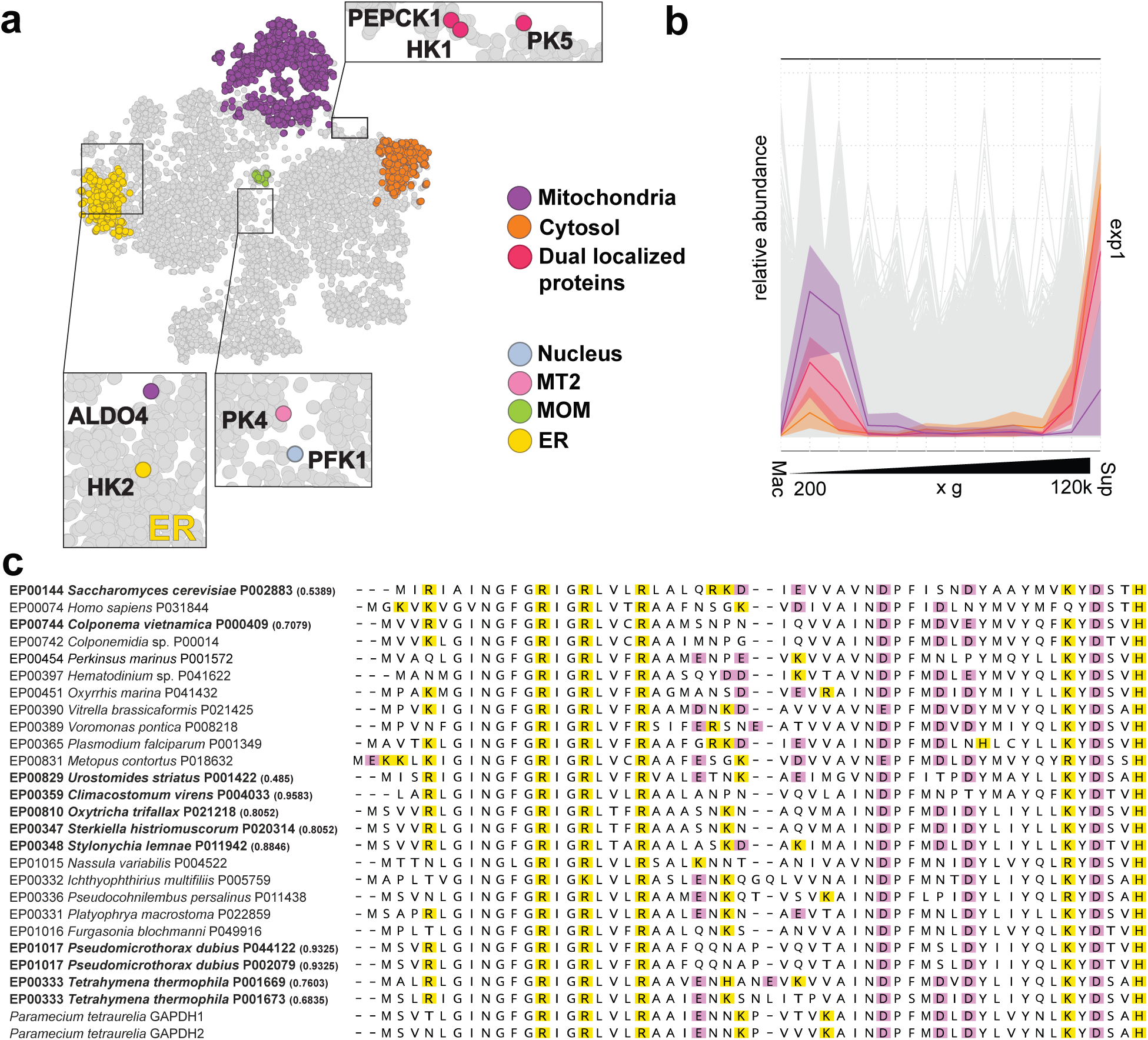
Dual localization of glycolytic and gluconeogenic proteins in Paramecium tetraurelia. (a) Some glycolytic and gluconeogenic proteins localize outside the mitochondrial and cytosolic clusters. PFK1, PK4 localize near the endomembrane and nuclear regions, whereas HK2, and ALDO1 localize near the ER cluster. PEPCK1, HK1, and PK5 localize in between the mitochondrial and cytosolic clusters. (b) PEPCK1, HK1, and PK5 exhibit protein abundance profiles intermediate to cytosolic and mitochondrial profiles indicating possible dual localization. (c) Highly conserved N-termini of GAPDH homologues in ciliates are often predicted as mitochondrial targeting signals suggesting relocalization of this enzyme to the mitochondria is possible in many organisms. Ciliate predicted proteomes were searched for GAPDH homologues and aligned using MUSCLE. Mitochondrial targeting signals were predicted using Target P.

Canonical mitochondrial targeting signals (MTSs) contain an N-terminal positively charged amphipathic helix ^68^. These MTSs are often, but not always, cleaved by the mitochondrial processing peptidase in the mitochondrial matrix ^69^. Though we often think of these targeting signals as definitive, dual localized proteins are well-known to exist ^70–72^, and some proteins can be targeted to mitochondria only under certain circumstances. Though a few of these examples are known from animals and other model systems, whether this phenomenon is widespread or generalizable is not known.

We used TargetP to determine which glycolytic or gluconeogenic enzymes carried N-terminal predicted MTSs. Because proteins used to train prediction tools are largely from animals, plants, or fungi, we used both the “plant” and “non-plant” options to predict MTSs. We found that the “plant” option usually gave higher mitochondrial targeting probabilities (Table S1) and therefore report these values in our figures and highlight values higher than 0.2. In *Paramecium*, of all the glycolytic and gluconeogentic enzymes, only TPI2-4 and PEPCK2-4 had MTS predictions scores above 0.2. Even though N-terminal sequences between the two ciliates didn’t greatly diverge (Figure 4C), *Tetrahymena*, TPI, GAPDH1-2, PGK, PGAM1-2, ENO2, PEPCK and PC all had MTS predictions higher than 0.2 (Table S1). These data suggest one of two possibilities. Either very small changes in an MTS (or its receptors) is enough for a complete relocalization of a protein, or certain proteins (e.g., glycolytic enzymes) may be targeted to different organelles in different conditions. Future investigations will uncover which possibility is more likely in ciliates.

## Conclusions

Subcellular proteomics has enabled us to predict over 4,500 proteins to 16 different subcellular compartments in *Paramecium tetraurelia*. Our data resolved several proteins to the MOM cluster highlighting several proteins that putatively function at ER-mitochondria contact sites. The glycolytic enzyme, PFK localized to the MOM cluster whereas several other glycolytic enzymes localized to the mitochondrial matrix. To explain these data, we suggest that ciliates favor gluconeogenesis over glycolysis and tend to use amino acids from digested proteins in both anabolic and catabolic carbon metabolism. We conclude that differential or possibly regulated retargeting of metabolic enzymes to different cellular compartments could be adaptive in either the physiological or evolutionary sense of the word. In the physiological sense, it is possible that glycolytic enzymes are relocated to different compartments based on the metabolic needs of the cell. In certain circumstances glycolytic enzymes may reside in the mitochondria, whereas at other times the proteins are in the cytosol. The cell is responding to its environment. In the evolutionary sense, it is possible that relocalizing parts of a metabolic cycle to different organelles provides a fitness advantage in particular lineages. Thus, in ciliates it may be beneficial to exhibit the mosaic distribution of glycolysis/gluconeogenesis, whereas in animals it may be beneficial to retain the canonical localizations. Future work will test each of these possibilities in turn.

## Acknowledgements

This work was funded by National Science Foundation grants (1927159, 2119963, and 2405455), Gordon and Betty Moore Foundation grant (GBMF10600), National Institute of Health grant 2R35GM122566, and VEDA FELLOWSHIPS within the Operational program Jan Amos Komensky (OP JAK), Project No. CZ.02.01.01/00/22_010/0008117.

## Author contributions

Conceptualization, methodology, and formal analysis, D.J., T.J.L, M.L., and J.G.W.; investigation, D.J., T.J.L, Y-P.P., N.Q., T.K., Y.Q., T-F.C., and J.G.W.; writing – original draft, T.J.L., D.J., and J.G.W.; writing – review & editing, T.J.L, D.J., Y-P.P., M.L., and J.G.W.

## Supplement Figure Legends

**Figure S1. Western blots of Paramecium fractions.** To confirm that different proteins are present with differing abundance across our fractions, we used immunoblotting using an anti-H3 antibody (NB500, abcam). 10 µg of protein was loaded into each lane. Detection was performed using an ECL Western Blotting Substrate and imaged using an Azure 600 (Biosystems). Image contrast was uniformly adjusted to enhance visibility.

**Figure S2. Paramecium subcellular proteome marker sets and classifications.** (a) Markers used for this investigation were plotted onto the t-sne. (b) SVM classification of all detected proteins to one of 16 compartments. (c) K-means clustering into 16 clusters. (d) Comparison of SVM clusters with k-means clusters.

**Figure S3. Phylogenetic reconstruction of alveolate dynamin related proteins.** Dynamin related proteins were collected from diverse alveolates from the EukProt database using a BLAST evalue cutoff of 10^-20^. Protein sequences were aligned using MUSCLE and trimmed manually. A phylogenetic tree was calculated using IQTREE.

**Figure S4. Tetrahymena thermophila subcellular proteome.** (a) Subcellular proteomics workflow. We used nitrogen cavitation to lyse harvested cells and followed the differential centrifugation protocol published by Geladaki et al. (2019). (b) Protein abundance profiles of mitochondrial, cytosol, mucocyst, and ribosomal clusters. After label-free quantification mass spectrometry, data processing, and mapping of marker proteins, we created abundance-distribution profiles of selected subcellular marker proteins measured in each experiment. Individual fractions contained different enrichments of organellar proteins. (c) To confirm that different proteins are present with differing abundance across our fractions, we used SDS-PAGE followed by staining with Coomassie brilliant blue. 10 µg of protein was loaded into each lane. Imaging was performed using an Azure 600 (Biosystems). Image contrast was uniformly adjusted to enhance visibility. (d) Subcellular protein atlas of *Paramecium tetraurelia*. We used t-SNE projection to visualize the distribution of organellar proteins after SVM classification. (e) Glycolytic and gluconeogenic proteins largely map to either the cytosol or the mitochondria in *Tetrahymena thermophila*.

**Figure S5. Tetrahymena thermophila supplement.** (a) Markers used for this investigation were plotted onto the t-sne. (b) SVM classification of all detected proteins to one of 13 compartments. (c) K-means clustering into 13 clusters. (d) Comparison of SVM clusters with k-means clusters.

**Figure S6. Location of TCA cycle, glycogenolysis, and related enzymes in ciliates.** Relevant TCA cycle enzymes were visualized in (a) *Paramecium* and (b) *Tetrahymena*. Relevant glycogenolysis enzymes were visualized in (c) *Paramecium* and (d) *Tetrahymena*.

**Figure S7. Tetrahymena growth curve in media containing different amounts of sugars.** The growth of ***Tetrahymena*** was monitored in four types of media with different glucose-to-protein ratios. Neff media was used as a control, SSP media was used to represent high-protein/low-glucose conditions, glucose-free modified Neff media for a protein-rich environment, and 2% glucose as a protein-free environment. The growth was monitored over a 5-day period at room temperature, and cell density was measured at 6-hour and 24-hour intervals using OD600. This procedure was carried out in biological triplicate.

**Table S1.** All protein ID tables in separate excel tabs.

## Resource availability

### Lead contact

Further information and requests for resources and reagents should be directed to and will be fulfilled by the lead contact, Jeremy Wideman.

### Materials availability

This study did not generate new materials.

### Data and code availability

Raw proteomic data were deposited on PRIDE PXD061592 (for *Paramecium tetraurelia*) and PXD061102 (for *Tetrahymena thermophila*).

Additional information required to reanalyze the data will be supplied upon request to the lead contact.

## Methods

### Culturing

#### Paramecium tetraurelia

Cultures of *Paramecium tetraurelia* strain 51 were a generous gift from Sascha Krenek (German Federal Institute of Hydrology). Cells were cultured using modified husbandry techniques described in Sonneborn ^13^. Approximately ∼1000 *P. tetraurelia* cells were inoculated into fresh wheat-grass medium (0.25% wheat grass powder, 0.05% Na_2_HPO_4_, Cerophyl, yeast extract, stigmasterol), which is bacterialized with stationary-phase *Klebsiella pneumoniae* to induce autogamy. DAPI staining was used to assess the macronuclear (MAC) state of each culture, scoring autogamous cells as those with fragmented MACs, and vegetative cells as those with intact MACs. When the culture contained ∼100% vegetative cells and the cell density reached ∼ 1000 cells/mL, the culture was harvested.

#### Tetrahymena thermophila

*Tetrahymena thermophila* SB 210 was obtained from *Tetrahymena* Stock Center (Cornell University, Ithaca, NY, USA). The initial culture was established and maintained in the modified Neff medium (0.25% proteose peptone, 0.25% yeast extract, 0.5% glucose, 33.3 mM FeCl_3_ • 6 H_2_O) at room temperature by serial passage every week as the lab stock. On the day before protein extraction, a 100 ml of a new culture was started from the lab stock (OD_600_ 0.9-1.1, ∼ 10^6^/mL) with 1:10 inoculation ratio and grown in SPP media (2% proteose peptone, 0.1% yeast extract, 0.2% glucose, 33.3 mM FeCl_3 •_ 6 H_2_O) at 30°C for 18-20 hrs with a constant agitation at 250 rpm.

### Cell harvesting

#### Paramecium tetraurelia

Vegetative *P. tetraurelia* cells were first sieved through a cheesecloth to remove bacterial biofilm and debris and then harvested using a 10μm nylon mesh. Cells were washed on the mesh with Dryl’s Solution (2 mM sodium citrate, 1 mM NaH_2_PO_4_, 1 mM Na_2_HPO_4_, 1mM CaCl_2_) and spun at 1000 × g for 10 min. Cell pellets were washed with detergent-free buffer (DF: 0.25 M sucrose, 10 mM HEPES pH 7.4, 2 mM EDTA, 2 mM Mg(OAc), Halt^TM^ Protease and Phosphatase Inhibitor Cocktail) (PPIs) and spun at 1000 × g for 5 mins, this step was repeated three times.

#### Macronuclear enrichment

Cell pellets for MAC isolation were resuspended in detergent-present lysis buffer (DP: 1% Triton-X, 0.25 M sucrose, 10 mM Tris, 3 mM CaCl₂, 8 mM MgCl₂, PPIs) and then dounced with 20–30 strokes to achieve 100% lysis efficiency. The cell homogenate was centrifuged at 300 × g for 5 mins and washed with DP buffer. This step was repeated three times, as were the subsequent washes and spins at 300 × g for 5 min with DF buffer. Prior to the final spin, an aliquot was stained with DAPI to visualize MAC enrichment by fluorescence microscopy. The MAC pellet was then stored at -80C.

#### Tetrahymena thermophila

Overnight cultures were harvested by centrifugation at 2,000×g for 2 minutes and then washed with 5 mL of 1X PBS. This step was repeated twice. After washing, cells were either lysed or subjected to the dibucaine assay that amputates the cilia and triggers the secretion of mature mucocysts ^73^. For the assay, cells were resuspended in 2 mL of assay buffer (10 mM HEPES, pH 7.0, 0.5 mM CaCl₂) and mucocyst excretion was stimulated for ∼30 seconds by adding 25 mM dibucaine dissolved in DMSO to final concentration 2.5 mM and gently inverting the tubes. After dibucaine exposure, cells were diluted to a 20 mL volume of assay buffer and pelleted at 1,000×g for 5 minutes. The released mucocyst content, which formed a flocculent layer above the pelleted cells, was removed using a serological pipette and discarded.

#### Grow curve

The growth of *Tetrahymena* was monitored in four types of media with different glucose-to-protein ratios. We used modified Neff media (0.25% proteose peptone, 0.25% yeast extract, 0.5% glucose, 33.3 mM FeCl_3 •_ 6 H_2_O) as a control, SSP media (2% proteose peptone, 0.1% yeast extract, 0.2% glucose, 33.3 mM FeCl_3 •_ 6 H_2_O) ^74^ to represent high-protein/low-glucose conditions, glucose-free modified Neff media for a protein-rich environment, and 2% glucose as a protein-free environment. A total of 60 mL of each media was placed into new flasks and inoculated with 1×10⁶ cells cultured in modified Neff media for 72 hours. The growth was monitored over a 5-day period at room temperature, and cell density was measured at 6-hour and 24-hour intervals using OD600. This procedure was carried out in biological triplicates for all media.

### Cell lysis and subcellular fractionation

#### Paramecium tetraurelia

The cells intended for a total cell lyses using nitrogen cavitation were resuspended in 6 mL of DF Buffer pre-chilled to 4°C and supplemented with PPIs. Suspension was transferred into a cell disruption bomb (Parr 4639, Parr Instrument Co.) pre-chilled to 4°C and connected to the nitrogen gas pressure tank. The bomb was then charged and equilibrated at 250 psi on ice for 10 minutes. After equilibration, the cell homogenate was collected from the outlet valve at the rate ∼3 drops/s, folding the foam into the liquid, and incubated again at 250 psi for an additional 5 min (Simpson 2010). Cell lysate was then differentially centrifuged following previously published protocol ^21^. Pellets were re-spun and all residual supernatant was discarded. Dry pellets were stored at -80C as was the cytosol-enriched supernatant.

#### Tetrahymena thermophila

Mucus-free and deciliated cells were resuspended in DF buffer and centrifuged at 1,000×g for 5 minutes. This step was repeated once more. The cell pellet was resuspended in 8 mL of pre-chilled DF buffer with PPIs. The cell suspension was divided into two equal parts for two different cell lysing protocols: douncing and nitrogen cavitation. Both methods break cells efficiently but gently, ensuring that all subcellular compartments remain intact, with an aim for ∼80–90% lysis efficiency. The 4 mL cell suspension was diluted to 8 mL with DF buffer and PPIs and transferred into a pre-chilled cell disruption bomb, bomb was equilibrated at 500 psi on ice for 10 minutes. After equilibration, the cell homogenate was collected from the outlet valve. A Dounce homogenization method was optimized using microscopic examination to assess the homogenate for cell breakage efficiency. 4 mL of the cell suspension in DF buffer and PPIs was transferred to a pre-chilled Teflon B-grade Dounce homogenizer. Cells were disrupted by 15 up-and-down strokes of a tight-fitting pestle. DAPI staining and microscopy was used to evaluate the cell lysis efficiency. The lysate was cleared by withdrawing the supernatant after centrifugation at 300×g for 5 minutes at 4°C to pellet unbroken cells, partially disrupted cells, and aggregates. The supernatant was transferred into a clean tube and kept on ice. The pellet was resuspended in 2 mL of DF buffer with PPIs, and the entire douncing procedure was repeated to achieve a sufficient amount of broken cells.Subcellular fractionation was performed by modifying the differential centrifugation protocol described previously ^21^. The centrifugation speeds and times were adjusted according to the protein composition and quantity visualized on the SDS-PAGE gel stained with Coomassie brilliant blue (Figure S4C).

### Protein distribution

#### Paramecium tetraurelia

We used immunoblotting to determine protein enrichment in individual *Paramecium* fractions using an anti-H3 antibody (NB500, abcam) (Figure S1). Pellets were resuspended in 2X Laemmli buffer (0.125 M Tris-HCl, pH 6.8; 4% SDS; 20% glycerol; 0.004% Bromophenol blue) without DTT. Samples were quantified using the Pierce™ BCA Protein Assay (Thermo Scientific), and 10 µg of protein was loaded onto an SDS-PAGE gel (Invitrogen™ Bolt™ Bis-Tris Plus Mini Protein Gels, 4-12%, 1.0 mm, WedgeWell™ format) along with a protein marker (Amersham™ ECL™ Rainbow™ Marker - Full Range). The gel was run for 1 hour at 110 V, briefly washed in 1X PBS buffer, and transferred onto a methanol-activated PVDF membrane (iBlot™ 2 Transfer Stacks, PVDF, Invitrogen™) using the iBlot™ 2 Western Blot Transfer Device (Invitrogen™). The membrane was blocked in 5% non-fat dry milk and 1X PBS buffer for 1 hour at room temperature, followed by overnight incubation at 4 °C with the anti-H3 antibody diluted 1:1000 in blocking solution (1% non-fat dry milk in 1X PBS buffer). The next day, the blots were washed three times for 10 minutes each in 1X PBS, probed with HRP-linked secondary antibodies (31460, invitrogen) diluted 1:1000 in blocking solution for 1 hour at room temperature, and rinsed again three times for 10 minutes each in 1X PBS-T. Detection was performed using the Pierce™ ECL Western Blotting Substrate (Thermo Scientific™), and imaging was conducted with the Azure 600 (Biosystems). Image contrast was uniformly reduced to enhance visibility.

#### Tetrahymena thermophila

Similarly to *Paramecium* samples, *Tetrahymena* fractions were quantified using the Pierce™ BCA Protein Assay (Thermo Scientific™), separated by SDS-PAGE gel (Bolt™ Bis-Tris Plus Mini Protein Gels, 4-12%, 1.0 mm, WedgeWell™ format, Invitrogen™). Gels were stained for 1 h in a 0.25% Coomassie brilliant blue solution (Pierce™ Coomassie Brilliant Blue Dyes, R-250, Thermo Scientific™) in methanol/water/acetic acid (1:10), destained by diffusion in the same solution minus the dye and stored in ddH_2_O to clear the background. Gels were imaged using the Azure 600 (Biosystems).

### Sample preparation and LC-MS Analysis

This step of the protocol was identical for all *Paramecium* and *Tetrahymena* samples.

Native protein pellets obtained from differential centrifugation were digested and desalted following the protocol for the S-Trap™ Micro Column (ProtiFi, USA). Protein concentration was quantified using the BCA assay (Thermo Scientific, USA), while peptide concentration was measured using a fluorometric kit (Thermo Scientific, USA).

#### Liquid-chromatography tandem mass spectrometry

LC-MS/MS analyses of *Paramecium* peptides were performed at the Biosciences Mass Spectrometry Core Facility, Arizona State University. Data-dependent mass spectra were collected in positive mode using an Orbitrap Fusion Lumos mass spectrometer coupled with an UltiMate 3000 UHPLC (Thermo Scientific). Peptides were fractionated on an Easy-Spray LC column (50 cm × 75 μm ID, PepMap C18, 2 μm, 100 Å) with an upstream trap column. Each sample was analyzed in technical triplicate. LC-MS settings: electrospray potential 1.6 kV, ion transfer tube temperature 300°C, and the “Universal” peptide analysis method. Full MS scans (375–1500 m/z) were acquired at a resolution of 120,000 with 3-second cycle times. The RF lens was set to 30%, AGC to “Standard,” and monoisotopic peak determination included charge states 2–7. Dynamic exclusion was 60 seconds with a 10 ppm mass tolerance. MS/MS spectra were acquired in centroid mode with a quadrupole isolation window of 1.6 m/z and CID energy of 35%. Peptides were eluted over a 240-minute gradient at 0.25 µL/min using 2–80% acetonitrile/water: 0–3 min (2%), 3–75 min (2–15%), 75–180 min (15–30%), 180–220 min (30–35%), 220–225 min (35–80%), 225–240 min (80–5%).

LC-MS/MS analysis of the digested peptides were performed on an EASY-nLC 1200 (Thermo Fisher Scientific) coupled to an Orbitrap Eclipse Tribrid mass spectrometer (Thermo Fisher Scientific). Peptides were separated on an Aurora UHPLC column (25 cm × 75 µm, 1.6 µm C18, AUR2-25075C18A, Ion Opticks) with a flow rate of 0.35 µL/min for a total duration of 135 min ionized at 1.6 kV in the positive ion mode. The gradient was composed of 2% solvent B (5 min), 2-6% B (7.5 min), 6-25% B (82.5 min), 25–40% B (30 min), 40-98% B (1min) and 98% B (15min); solvent A: 2% ACN and 0.2% FA in water; solvent B: 80% ACN and 0.2% FA. MS1 scans were acquired at the resolution of 120,000 from 350 to 1,600 m/z, AGC target 1e6, and maximum injection time 50 ms. MS2 scans were acquired in the ion trap using fast scan rate on precursors with 2-7 charge states and quadrupole isolation mode (isolation window: 0.7 m/z) with higher-energy collisional dissociation (HCD, 30%) activation type. Dynamic exclusion was set to 30 s. The temperature of ion transfer tube was 300°C and the S-lens RF level was set to 30.

#### Tetrahymena thermophila

Tetrahymena samples were processed at the Proteome Exploration Laboratory, Beckman Institute, California Institute of Technology. LC-MS/MS analysis of the digested peptides were performed on an EASY-nLC 1200 (Thermo Fisher Scientific) coupled to an Orbitrap Eclipse Tribrid mass spectrometer (Thermo Fisher Scientific). Peptides were separated on an Aurora UHPLC column (25 cm × 75 µm, 1.6 µm C18, AUR2-25075C18A, Ion Opticks) with a flow rate of 0.35 µL/min for a total duration of 135 min ionized at 1.6 kV in the positive ion mode. The gradient was composed of 2% solvent B (5 min), 2-6% B (7.5 min), 6-25% B (82.5 min), 25–40% B (30 min), 40-98% B (1min) and 98% B (15min); solvent A: 2% ACN and 0.2% FA in water; solvent B: 80% ACN and 0.2% FA. MS1 scans were acquired at the resolution of 120,000 from 350 to 1,600 m/z, AGC target 1e6, and maximum injection time 50 ms. MS2 scans were acquired in the ion trap using fast scan rate on precursors with 2-7 charge states and quadrupole isolation mode (isolation window: 0.7 m/z) with higher-energy collisional dissociation (HCD, 30%) activation type. Dynamic exclusion was set to 30 s. The temperature of ion transfer tube was 300°C and the S-lens RF level was set to 30.

### Raw data processing and quantification

#### Paramecium tetraurelia

Label-free quantification (LFQ) was performed using Proteome Discoverer 2.4 (Thermo Scientific) based on the composite database: *P. tetraurelia* strain 51’s predicted proteome (https://paramecium.i2bc.paris-saclay.fr/), 31 mitochondrial ORFs, *K. aerogenes* predicted proteome (https://www.uniprot.org/), and common laboratory contaminants (cRAP; https://www.thegpm.org/crap/). Raw files were searched with SequestHT using Trypsin as the enzyme, allowing up to three missed cleavages. Peptide length was set to 6–144 amino acids, with precursor ion mass tolerance at 20 ppm, fragment mass tolerance at 0.5 Da, and a minimum of one peptide identified. Carbamidomethyl (C) was a fixed modification, while Acetyl (N-terminus), Met-loss (N-terminus), and oxidation of Met were dynamic modifications. A target/decoy strategy and 1.0% FDR were calculated using Percolator. Data were imported into Proteome Discoverer 2.4, and features were detected using the Minora Feature Detector algorithm. The area-under-the-curve (AUC) for aligned ion chromatograms was calculated to determine relative abundances. The RAW data have been deposited to the ProteomeXchange Consortium via the PRIDE partner repository with the dataset identifier PXD061592.

Proteins and their corresponding LFQ abundance values were imported into the R programming language and converting into MSnset object using the Bioconductor packages MSnbase (v 2.20.1) and pRoloc (v 1.34.0) ^27^. The data were checked and proteins from cRAP or *K. aerogenes*, or with low confidence (PSM < 3 and without uni-peptide) were filtered out. The proteins identified only in the MAC or Sup fractions were also excluded from the following analysis. Technical triplicates were averaged to generate a 36th dimensional dataset of relative protein abundance. The datasets were split into their respective experiments (i.e., 1-12, 13-24, 25-36) to perform hybrid imputation and sum-normalization across rows.

Missing data were imputed first by nearest-neighbor averaging and then imputing zeros for all remaining empty cells. Principal component analysis and t-distributed Stochastic Neighbor Embedding (tSNE) were applied for dimensional reduction and data visualization.

#### Tetrahymena thermophila

Raw data files were analyzed using Proteome Discoverer (PD) 2.5, Thermo Fisher Scientific) based on the CHIMERYS or Sequest algorithm against in silico tryptic digested Tetrahymena reference proteome database (https://tet.ciliate.org/downloads.php). The maximum missed cleavages were set to 2. Dynamic modifications were set to oxidation on methionine (M, +15.995 Da) and carbamidomethylation on cysteine residues (C, +57.021 Da) was set as a fixed modification. The maximum parental mass error was set to 10 ppm, and the MS2 mass tolerance was set to 0.3 Da. The false discovery threshold was set strictly to 0.01 using the Percolator Node validated by q-value. The relative abundance of parental peptides was calculated by integration of the area under the curve of the MS1 peaks using the Minora LFQ node. The RAW data have been deposited to the ProteomeXchange Consortium via the PRIDE partner repository with the dataset identifier PXD061102. The resulted abundance profiles from Proteome Discoverer were imported into R ^75^ and analyzed with the packages Msnbase 2.8.3 and pRoloc 1.38.2 ^27,76^. The concatenated dataset from the three experiments yielded 7229 protein abundance profiles across 33 samples.

The data were further filtered by eliminating proteins with low confidence, low number of PSM (< 3) and without uni-peptide. One fraction, pellets from the 15k spin, of experiment 1, was excluded due to poor quality (missing data > 99%) resulting in data from 5680 proteins. Missing data were imputed first by nearest-neighbor averaging and then imputing zeros for all remaining empty cells. Principal component analysis and t-distributed Stochastic Neighbor Embedding (tSNE) were applied for dimensional reduction and data visualization.

### Supervised and Unsupervised Classification

#### Paramecium tetraurelia

For *Paramecium*, 291 manually curated marker proteins (Table S1) were used as the training set for a support vector machine (SVM) model with the svmOptimization and svmClassification functions in pRoloc package. Initially, 100 rounds of five-fold cross-validation were performed to optimize the SVM parameters based on the marker protein abundance profiles. The optimal parameters for the SVM classifier were then applied to all proteins in the dataset with a corresponding SVM score whose range is 0-1 with 1 being the score of marker proteins. The SVM classifier was then applied to unlabeled data (i.e., non-marker proteins) with corresponding weights applied to each marker class. Each protein was thus classified to one compartment, and any protein whose classification fell below the global median SVM score was reset to ‘unknown’ while the other half of the dataset was considered “predicted” to its corresponding compartment due to their higher SVM scores (Figures 1C and Table S1).

Unsupervised clustering was performed using the K-means (KM) algorithm implemented in the MLearn function from the MLInterfaces package in R (version 1.67.1). KM generated k random centroids and includes surrounding data points iteratively such that all data points are included in one of the k clusters and the size of each centroid is minimized. Kmeans clusters were generated with 16 clusters (Figure S2C). Comparison between kmeans clusters and svm clusters performed and plotted (Figure S2D).

#### Tetrahymena thermophila

For *Tetrahymena*, 187 manually curated marker proteins (Table S1), from 13 subcellular compartments, were used to classify all 5680 proteins. To classify all the proteins into the subcellular compartments, we first used supervised classification by support vector machine (SVM) model with the functions, svmOptimization and svmClassification, implemented in pRoloc. For all the clusters, median was used to set up the cut-off scores, and the proteins with SVM score greater than the cutoff was considered to successfully predict its corresponding compartment. The tSNE map of the SVM predicted proteins were plotted by ShinyApp implemented in pRolocVis 1.38.2. Unsupervised k-means algorithm implemented in the MLearn function from the MLInterfaces 1.78.0 in R to generate 13 K-means clusters, in which the proteins distribution profiles within groups are more alike than between groups.

### Targeting sequence identification and pathway analysis

The overexpression of *Paramecium* proteins after trichocyst discharge and ciliary shedding was also determined based on the RNA-seq profiles ^37^. MitoFates 1.2 ^35^ was used to predict the mitochondrial-targeting signals. Proteins with MTS score greater than 0.9 were considered as mitochondrial-targeting proteins. DeepTMHmm 1.0.24 ^36^ was applied to predict the transmembrane topology and SignalP 6.0 ^33^ was further used to identify if the signal peptide existed for Sec/SPI recognition. NLStradamus r.9 ^34^ was applied to predict the NLS existence and identify the proteins potentially processed by the nuclear import pathway. Metabolic pathway analyses were done with the KEGG pathway database (https://www.genome.jp/kegg/pathway.html).

